# β-cyclocitric acid: a new apocarotenoid eliciting drought tolerance in plants

**DOI:** 10.1101/478560

**Authors:** Stefano D’Alessandro, Yusuke Mizokami, Bertrand Legeret, Michel Havaux

## Abstract

β-Cyclocitral (β-CC) is a volatile compound deriving from ^1^O_2_ oxidation of β-carotene in plant leaves. β-CC elicits a retrograde signaling, modulating ^1^O_2_-responsive genes and enhancing tolerance to photooxidative stress. Here, we show that β-CC is largely converted into β-cyclocitric acid (β-CCA) in leaves and that this metabolite is a signal involved in stress tolerance. Treatment of Arabidopsis plants with β-CCA markedly enhanced plant tolerance to drought by a mechanism different from known responses such as stomatal closure, changes in osmotic potential and jasmonate signaling. Furthermore, we show that the response to β-CCA does not fully overlap with the β-CC-dependent signaling, indicating that β-CCA induces only a branch of the β-CC signaling pathway. In addition, the protective effect of β-CCA is a conserved mechanism, being observed in a variety of plant species. This study provides a new bioactive agent with promising agronomic applications for protecting plants against drought.

---

Oxidation of the carotenoid β-carotene by reactive oxygen species (ROS), especially singlet oxygen (^1^O_2_), produces various derivatives (apocarotenoids) including β-cyclocitral (β-CC) ^1,2^. This phenomenon was shown to take place in plant leaves and enhanced under stress conditions ^1,2^. In fact, when plants are exposed to environmental constraints (i.e. drought, cold or pathogens), which inhibit the photosynthetic activity, light energy can be absorbed in excess to what can be used by the photosynthetic processes, hence favoring transfer of electrons or excitation to oxygen and leading to ROS formation ^3,4^. Singlet oxygen is produced from triplet-excited chlorophylls, mainly in the PSII reaction centers ^5–8^. In fact, the PSII centers bind several β-carotene molecules that can scavenge ^1^O_2_ molecules generated therein ^5,9^. ^1^O_2_ quenching by carotenoids proceeds by a physical mechanism that leads to thermal energy dissipation ^10^ and also through a chemical quenching mechanism involving direct oxidation of the carotenoid molecule by ^1^O_2_ ^2,5,11^. Thus, as a major site of ^1^O_2_ production, PSII is also a major generator of oxidized β-carotene metabolites such as β-CC.

β-CC (SI Appendix, Fig. S1*A*) is a volatile compound that was shown to act as a signal molecule in Arabidopsis (*Arabidopsis thaliana*), triggering changes in the expression of ^1^O_2_-responsive genes and leading to acclimation to ^1^O_2_ and photooxidative stress ^1^. Produced by excessive light excitation at the level of PSII in the chloroplast, β-CC can be considered as an upstream mediator in the ^1^O_2_ retrograde pathway leading to acclimation. Actually, β-CC is one among several signaling metabolites that have been recently associated with chloroplast-to-nucleus retrograde signaling ^12–14^. However, for most of them including β-CC, the primary targets are still unknown. Here, we show that β-CC is converted to β-cyclocitric acid (β-CCA) not only in water as previously reported ^15^ but also *in vivo*, thus constituting one of the first step in the β-CC-dependent signaling. Exposing plants to exogenous β-CCA induces the expression of β-CC- and ^1^O_2_-responsive genes and enhances plant resistance to photooxidative conditions such as water stress. Since β-CCA is a water-soluble molecule that can be easily applied to plants, *e.g*. through irrigation water, those results support the possibility of using this compound to boost drought resistance of crops.

## Results and Discussion

β-CC can oxidize into β-cyclocitric acid (β-CCA, 2,2,6-trimethylcyclohexene-1-carboxylic acid), also known as β-cyclogeranic acid (SI Appendix, Fig. S1A, B). This conversion occurs spontaneously, *e.g.* upon addition of β-CC in water ^15^, as confirmed in Fig. S1C (SI Appendix). When injected in water, β-CC disappeared within 24 h, with the concomitant appearance of β-CCA as major oxidation product (SI Appendix, Fig. S1C) ^15^. The question arises as to whether oxidation of β-CC into β-CCA takes place *in vivo* too. Using GC-MS, we were able to measure β-CCA in control Arabidopsis leaves, and the measured concentrations were even higher than β-CC levels (Fig. 1*A*). This relative accumulation of β-CCA compared to β-CC was amplified under stress conditions: when plants were exposed to drought stress, the β-CC concentration rose by a factor of 3, revealing a condition of excessive light and photooxidative stress ^1,16^, while a 15-fold increase in β-CCA was observed (Fig. 1*B*). Moreover, when plants were treated for 4 h with volatile β-CC in a closed Plexiglas box (as previously described ^1^), the increased levels of β-CC in the leaves (about x 3) were found to be associated with a strong accumulation of β-CCA (Fig. 1*C*), showing that the conversion of β-CC into β-CCA does take place *in vivo.* The oxidation of β-CC in water without adding any oxidizing reagent (besides dissolved O2) was suggested by Tomita et al. ^15^ to proceed according to the Baeyer-Villiger oxidation mechanism which produces esters from ketones and carboxylic acids from aldehydes ^17^. This β-CC-to-β-CCA conversion in Arabidopsis appeared to occur with a very high efficiency since the accumulation levels of β-CCA in β-CC-treated plants were much higher than the β-CC accumulation levels (Fig. 1*C*). Therefore, we cannot exclude that β-CCA formation is facilitated by an enzyme-catalyzed reaction *in planta, e.g.* by a Baeyer-Villiger monooxygenase ^18^, as previously reported for the oxidation of castasterone to brassinolide in brassinosteroid biosynthesis ^19^.

**Fig. 1.**
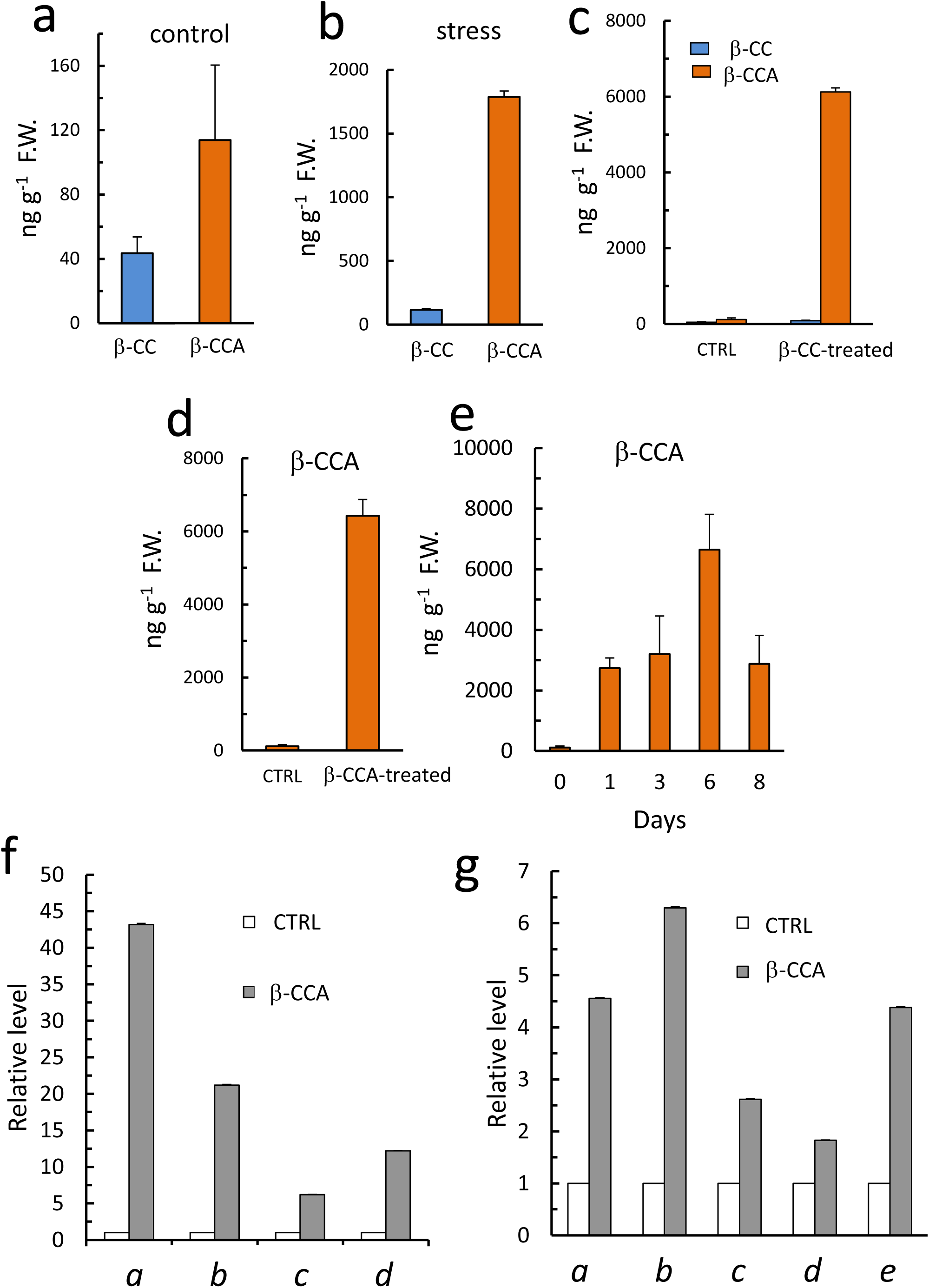
β-CCA levels in leaves of Arabidopsis plants and their effect on gene expression. a) Control, untreated plants (5 plants). b) Plants exposed to water stress (no watering for 7 d, 2 plants). c) Plants treated for 4 h with 100 μl volatile β-cc in a hermetically closed box (2 plants). The controls were treated similarly with 100 μl water. d) β-CCA levels in Arabidopsis leaves sprayed with 50 μl/leaf of 1 mM β-CCA or of water (control). Leaves were taken 24 h after the treatment (2 plants). e) β-CCA levels in leaves of Arabidopsis plants watered at time 0 with 25 ml of a 1 mM β-CCA solution or with watered acidified with 1 mM citric acid (6 plants pooled, 2 technical replicates). f) Expression levels of ^1^O_2_- and β-CC-responsive genes analyzed by qRT-PCR in control leaves and in leaves sprayed with β-CCA (3 plants pooled, 3 technical replicates). g) Expression levels of ^1^O_2_- and β-CC-responsive genes analyzed by qRT-PCR in control plants and in plants watered with β-CCA or with 1 mM citric acid for 24 h (3 plants pooled, 3 technical replicates). In panels f and g, *a*= AT3G04000, *b*= AT5G61820, *c*= AT5G63790, *d*= AT5G16970, *e*= AT3G28580.

When attached leaves were directly sprayed with β-CCA (Fig. 1*D*), a strong accumulation of β-CCA was measured inside leaf tissues, indicating that this compound can readily enter the leaves. Since β-CCA is soluble and stable in water contrary to β-CC, it can be easily applied to whole plants through irrigation. In Fig. 1*E*, Arabidopsis plants, treated with β-CCA through the soil, showed β-CCA accumulation in the leaves. This indicates that exogenously applied β-CCA is taken up by the roots and transported to the leaves through the xylem. Whether applied through roots or directly on leaves, β-CCA accumulated in leaves without any significant change in the β-CC content (SI Appendix, Fig. S2). Since β-CCA is directly formed from β-CC, *de novo* synthesis of β-CCA would imply an increased synthesis of β-CC which was not observed, making improbable that the β-CCA treatment triggered β-CCA synthesis rather than β-CCA fluxes from the soil to the plant tissues.

The expression of a number of genes that were previously identified as responsive to β-CC ^1^ was analyzed by qRT-PCR before and after application of β-CCA by spraying attached leaves or by watering plants: AT3G04000 (*ChlADR*), AT5G61820 (unknown), AT5G63790 (*ANAC102*), AT5G16970 (*ALKENAL REDUCTASE AER*), AT3G28580 (*AAA+ ATPASE*). Both direct application of β-CCA on leaves and watering of plants with a solution of β-CCA induced a strong upregulation of the selected genes (Fig. 1F and G), indicating that β-CCA acts as a signaling molecule triggering transcriptomic changes. The gene up-regulation levels were lower in plants watered with β-CCA relative to leaves directly sprayed with β-CCA, as were the accumulation levels of β-CCA inside the leaf tissues (24 h after treatment).

A previous microarray-based transcriptomic study revealed that a number of water stress-responsive genes are also inducible by β-CC ^1^. The same phenomenon was observed for β-CCA by qRT-PCR analyses (Fig. 2*A*): the drought marker genes *ATAF1* (ANAC002, AT1G01720), *RD29A* (AT5G52310), *RD29B* (AT5G51180), *RD22* (AT5G25610), *bZip60* (AT1G42990) ^20–23^ were induced by β-CCA in the absence of any water stress. However, not all drought marker genes responded to β-CCA since *DREB2A* (AT5G05410), a transcription factor functioning in water stress response ^24^, was not induced. *RD26* (*ANAC072*, AT4G27410) and *bZIP60* (AT1G42990) showed a different response to β-CCA compared to the other genes, since their induction was transient. Nevertheless, our results indicate that the response elicited by β-CCA overlaps, at least partially, with the genetic response to water stress. This response may reflect the role of ^1^O_2_ in water stress ^25^. This prompted us to analyze the effect of β-CCA on Arabidopsis plants exposed to drought stress. Following irrigation of Arabidopsis plants with water containing or not β-CCA (same pH), water stress was induced by withdrawing irrigation. After 7 d, control plants showed clear symptoms of stress and dehydration (Fig. 2*B*). These symptoms were strongly attenuated in plants pretreated with β-CCA. After rewatering with plain water, control plants did not recover and died, whereas β-CCA-treated plants recovered and were fully turgescent (Fig. 2*C*). The leaf relative water content (RWC) was much higher in plants treated with β-CCA throughout the water stress treatment compared to control plants (Fig. 2*G*). The protection of Arabidopsis by β-CCA against drought stress was confirmed in other plant species such as pepper (*Capsicum*), pansy flower plants (*Viola tricolor*) (SI Appendix, Fig. S3) and tomato (*Solanum lycopersicum*, see below).

**Fig. 2.**
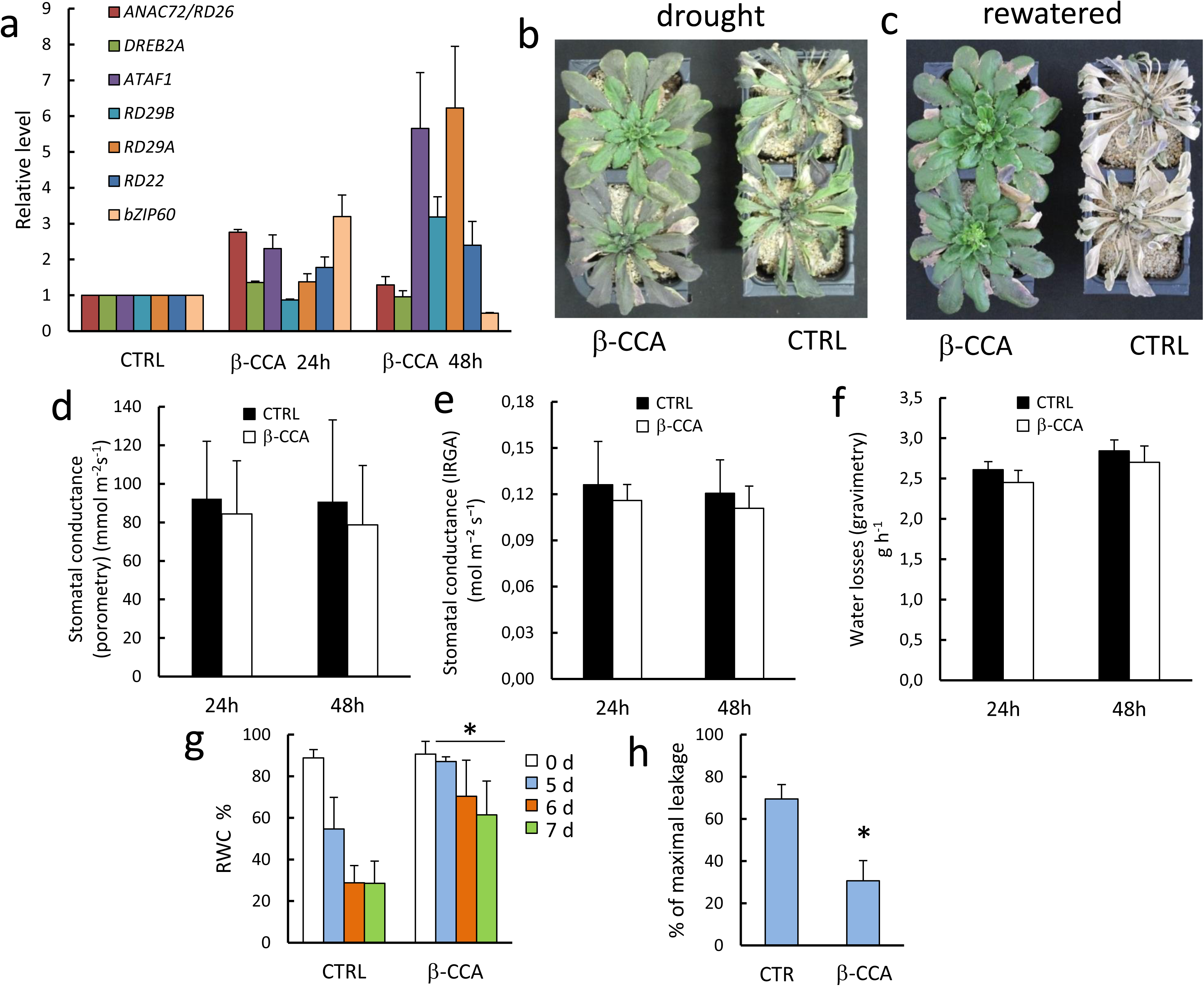
β-CCA-induced protection of Arabidopsis plants against drought stress. a) Expression levels of water stress-responsive genes in leaves of Arabidopsis plants watered with β-CCA or with citric acid (3 plants pooled, 4 technical replicates). b) Picture of Arabidopsis plants pre-treated with β-CCA or with citric acid (CTRL) and then subjected for 7 d to drought stress imposed by withdrawing watering (2 plants). c) Picture of the same plants 10 d after re-watering with plain water. d-e) Stomatal conductance measured by porometry (2 plants, mean value of 10 leaves per point) (d) or by IRGA analyses of gas exchange (5 plants) (e). f) Losses of water by Arabidopsis plants (treated or untreated with β-CCA) during water stress (3 plants). g) RWC of plants subjected to water stress after pre-treatment with β-CCA or with acidified water (2 plants, mean values of 6 leaves per point). h) Increased membrane permeability in Arabidopsis leaves, as measured by electrolyte leakage, after 7 d of water stress (3 plants, 6 leaves were pooled). *, different from CTRL at P <0.01 (Student’s t-test).

Typical responses of plants to water stress are stomatal closure mediated by the apocarotenoid abscisic acid, hence reducing transpiration and water losses ^26–28^, osmotic adjustment ^29,30^ and jasmonate signaling ^31^. Although, β-CCA is an apocarotenoid as well as abscisic acid, the protective role of β-carotene against drought stress does not rely on stomatal regulation. In fact, stomatal closure was not induced by β-CCA (Fig. 2*D* and *E*), and transpiration was similar in β-CCA-treated and -untreated plants (Fig. 2*F*). We also examined the effects of β-CCA on the *ost2-2* Arabidopsis mutant that is impaired in stomatal regulation due to a constitutive activation of a plasma membrane ATPase ^32^ and on the *abi1* mutant affected in ABA signaling ^33^. As expected, stomatal conductance and transpiration of *ost2-2* leaves were enhanced compared to WT leaves (Fig. 3*C*), and water stress developed much more rapidly in *ost2-2* plants relative to WT plants when water irrigation was stopped. However, the protective action of β-CCA was confirmed in the mutant: leaf dehydration and loss of turgescence were less pronounced in β-CCA-treated *ost2-2* plants (Fig. 3*A* and *B*). As with WT plants (Fig. 2), the RWC of *ost2-2* mutant plants was preserved by β-CCA during water stress (Fig. 3*B*). Similar to *ost2-2*, the tolerance of the *abi1* mutant to drought was noticeably increased by β-CCA (Fig. 3*E* and *F*). The protection of *ost2-2* and *abi1* by β-CCA confirms that the mode of action of β-CCA does not rely on ABA signaling and the associated stomatal closure. Also, plant response to water stress usually involves synthesis of osmoprotectants, such as ammonium compounds, sugar, sugar alcohols and amino acids ^29,30^, which permit the maintenance of turgor pressure under water stress conditions, thus preserving vital functions. We checked whether accumulation of β-CCA in leaves could bring about substantial changes in leaf osmotic potential Ψ_π_. However, the data of Fig. 4A show that the protective function of β-CCA against drought stress does not rely on this phenomenon. Indeed, Ψ_π_ (−0.815 MPa) of leaves taken from plants watered with β-CCA did not differ significantly from Ψ_π_ of control leaves (−0.786 MPa). Finally, the involvement of a jasmonic acid dependent mechanism ^31^ in the β-CCA effect was excluded. In fact, the Arabidopsis mutant *coi1,* that lacks the jasmonate receptor COI1, although more sensitive than the WT, responded to β-CCA like the WT, by an increase in drought tolerance (Fig. 4B and C).

**Fig. 3.**
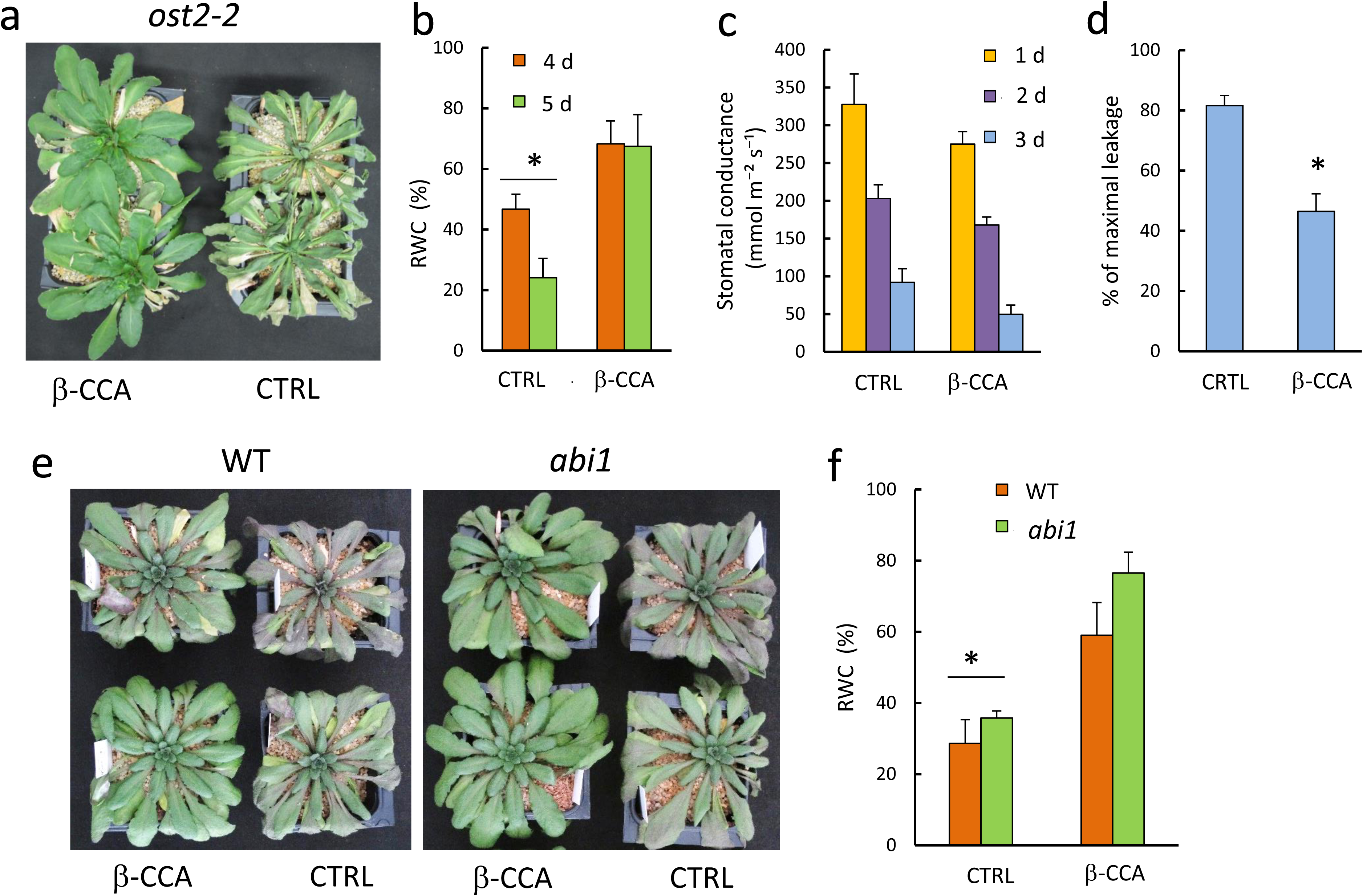
β-CCA-induced protection of Arabidopsis *ost2* and *abi1* mutant plants against drought stress. (a) Picture of *ost2-2* mutant plants pre-treated with β-CCA or with water (CTRL) and then subjected to 5 days drought stress (2 plants). (b) and (c) Stomatal conductance (2 plants, mean value of 10 leaves per point) and RWC (2 plants, mean values of 6 leaves per point) of WT or *ost2-2* plants pre-treated with β-CCA or with water and then subjected to water stress. (d) Membrane permeability in *ost2-2* mutant leaves measured by ion leakage after 5 d of water stress (2 plants, 6 leaves were pooled). (e) Picture of *abi1* mutant plants pre-treated with β-CCA or with water (CTRL) and then subjected to 5 days drought stress (2 plants). (f) RWC of leaves (2 plants, mean values of 6 leaves per point). *, different from CTRL at P <0.01 (Student’s t-test).

**Fig. 4.**
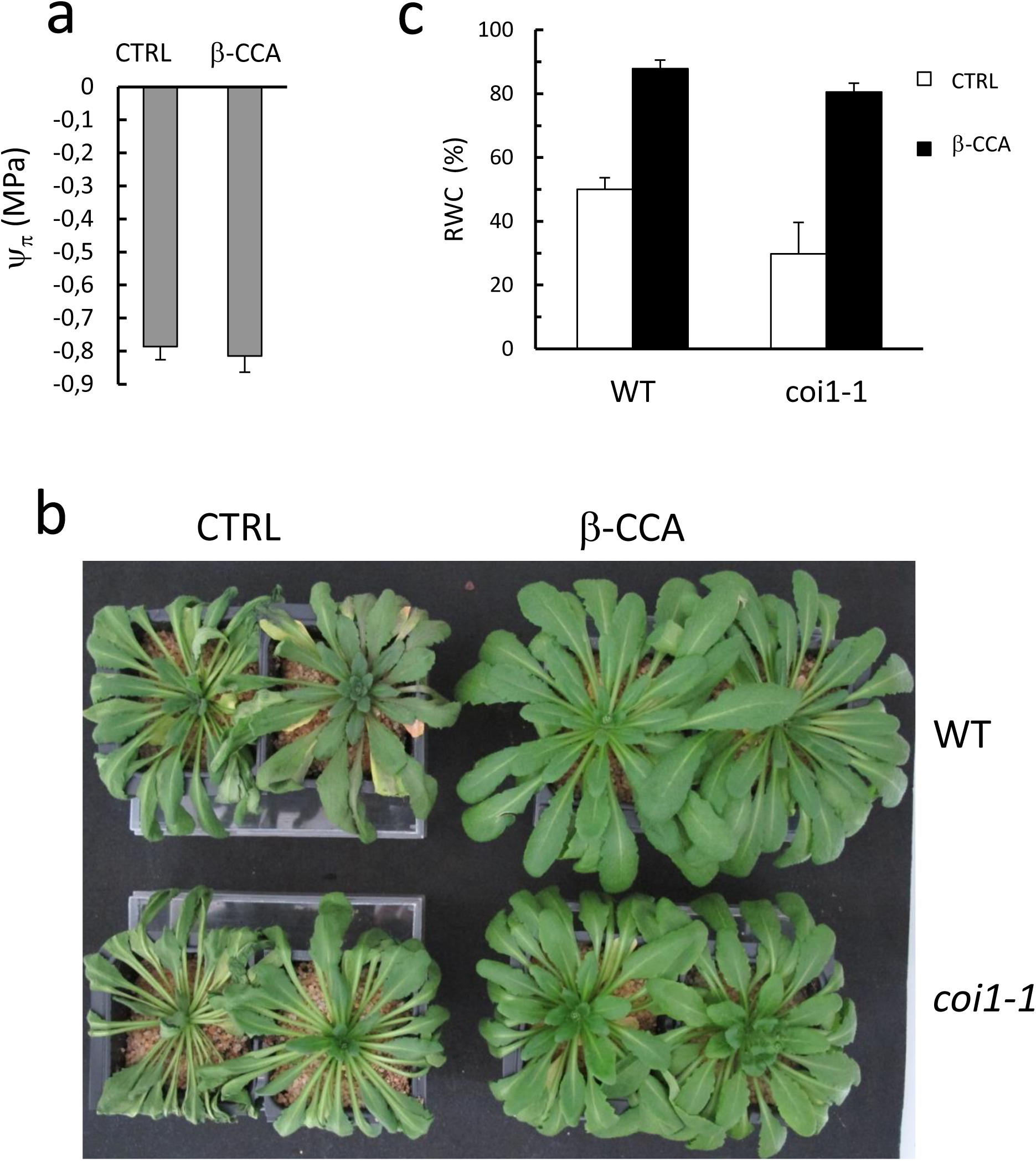
Neither osmotic adjustment, nor jasmonate signaling are required for the β-CCA protective effect. a) Leaf osmotic potential of Arabidopsis plants watered with 1 mM β-CCA for 48h. Ctrl, control (8 plants). b). Effect of water deprivation on WT Arabidopsis plants and on *coi1* mutant plants pre-treated with 0 or 1 mM β-CCA (2 plants). c) RWC of the leaves (2 plants, mean values of 6 leaves per point).

β-CCA protected cell membranes against damage under drought stress, as indicated by a much lower leakage of electrolytes by leaves from β-CCA-treated plants compared to untreated plants, in both the WT and *ost2-2* backgrounds (Fig. 2*H*, Fig. 3*D*). It is thus possible that β-CCA induces a cellular defense mechanism that reinforces membrane stability and resistance to ROS. It is likely that this protection against membrane disruption during water stress contributed to maintaining leaf water content and enhancing drought tolerance ^34–36^. The preservation of cell membrane stability is anyway an effect recalling β-cc induced acclimation to photooxidative stress ^1^. Unfortunately, the β-CC-dependent signaling pathway is still largely unknown. Nevertheless, the role of *METHYLENE BLUE SENSITIVITY 1* (*MBS1*) ^37^ and of the SCARECROW LIKE 14 (SCL14)-dependent detoxification pathway in the response to β-cc have been recently described ^16,38^. MBS1 is a cytosolic zinc finger protein essential for the regulation of the expression of ^1^O_2_-responsive genes ^37^. In the *mbs1* mutant that lacks the MBS1 protein, the expression of ^1^O_2_ marker genes was shown to be markedly deregulated ^37^. For some genes, induction by β-CC was blocked while the induction of other genes was enhanced ^16^. When exposed to drought stress, the *mbs1* mutant behaved as WT, with β-CCA providing a marked protection against water stress deprivation (Fig. 5 *A* and *B*). We can therefore conclude that the MBS1-dependent signaling pathway, triggered by β-CC and essential for the β-CC induced acclimation to photooxidative stress, is different from the β-CCA-induced signaling pathway leading to drought stress resistance. Consistently, drought-related genes whose expression was observed to be upregulated by β-CCA (Fig. 2*A*) were also induced in the *mbs1* mutant (Fig. 5*C*). It is therefore possible that β-CCA mediates a branch of the β-CC signaling pathway while MBS1 regulates another (or several other) branch(es).

**Fig. 5.**
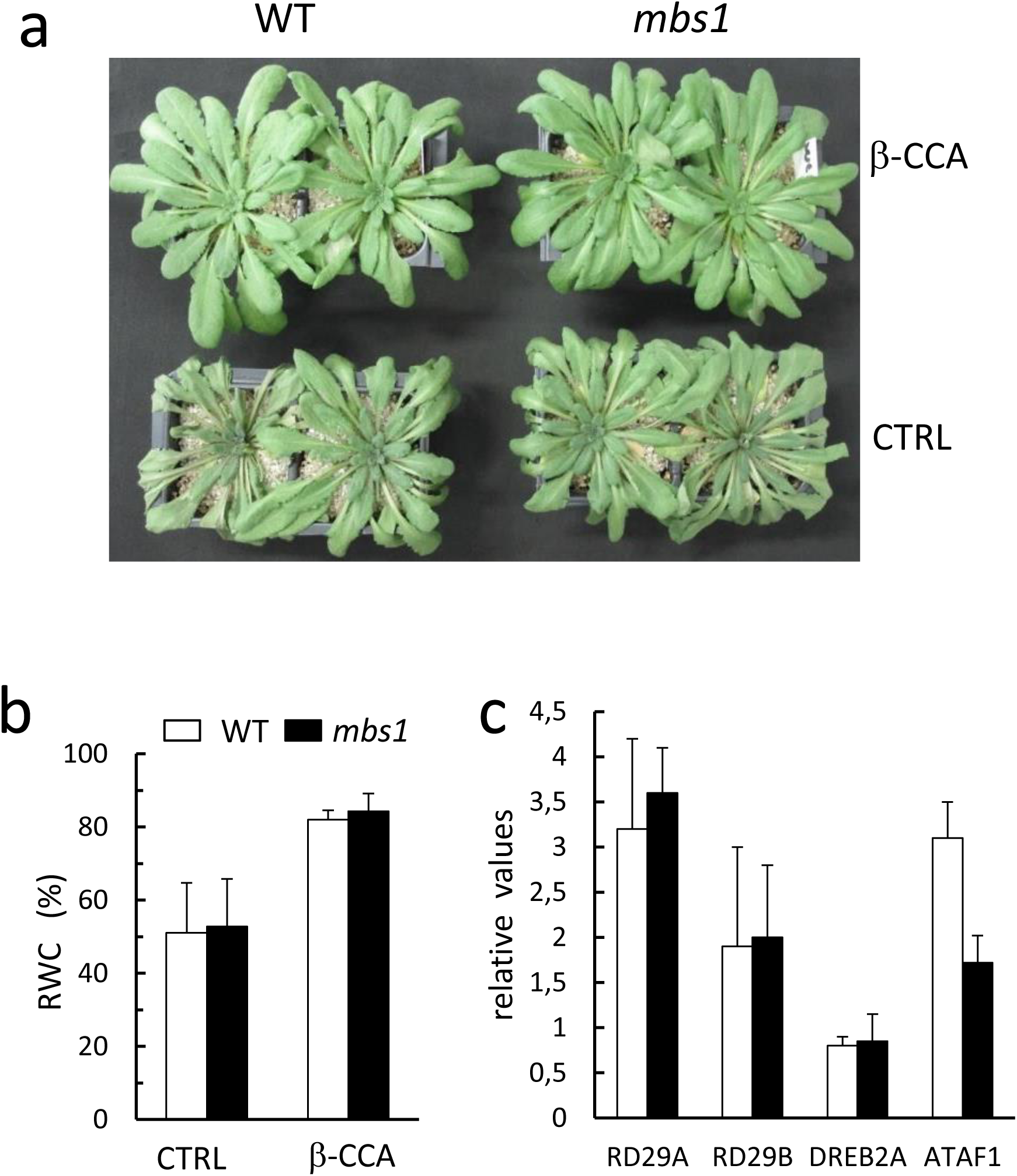
The *mbs1* mutation does not suppress the protective effect of β-CCA against drought stress. a) Picture of the plants (WT and mbs1 mutant) pre-treated with β-CCA or with water (CTRL) and exposed to water stress for 7 d (2 plants). b) RWC of WT and mbs1 leaves after 7 days of water stress (2 plants, mean values of 6 leaves per point). c) Expression levels of several drought-responsive genes (3 plants pooled, technical 4 replicates).

Moreover, in contrast with β-CC ^1^, β-CCA did not provide any protection against photooxidative stress induced by excessive light (SI Appendix, Fig. S4) which have been previously shown to depend on MBS1 and on the SCL14-dependent detoxification pathway ^16,38^. We, therefore, tested the involvement of the more recently described SCL14-dependent branch of the β-CC response ^38^, by the use of the *scl14* knockout mutant and of the OE:SCL14 overexpressing line. While the mutant line showed no difference with the WT in the response to the β-CCA treatment (data not shown), the SCL14 overexpressing line did show an enhanced resistance to drought stress already when treated with water, and even stronger drought tolerance when treated with β-CCA (Fig. 6A and B). In fact, the RWC of WT plants treated with β-CCA and of OE:SCL14 plants treated with water or with β-CCA after 7 days of water withdrawal was markedly higher (70 – 90 % RWC) than WT plants treated with water (20 % RWC). Furthermore, the overexpression of SCL14 and the treatment with β-CCA had an additive effect, since OE:SCL14 plants treated with β-CCA showed an RWC of 60% after 11 days of water withdrawal, indicating an almost doubled resistance compared to WT plants treated with water. SCL14 is a GIBBERELLIN-ACID INSENSITIVE (GAI), REPRESSOR of GA1 (RGA) and SCARECROW (SCR) (GRAS) protein involved in the regulation of the xenobiotic detoxification response ^39^ and more recently described in the response to photooxidative stress under excessive light ^38^. We, thus, quantified the expression levels of four reporter genes in the SCL14-dependent response: *ANAC102, ATAF1, SDR1* and *ChlADR*, by qRT-PCR ^38,39^. The four reporters were slightly induced at 24h of treatment with β-CCA, to levels similar to the ones found in the OE:SCL14, and only the treatment of the SCL14 overexpressing lines with β-CCA generated a marked induction of the detoxification pathway (Fig. 6C). The weak induction of this pathway by a 24-hour treatment with β-CCA is in line with the lack of protection of this treatment to excessive light conditions (SI Appendix, Fig. S4). At the same time, we cannot exclude that this pathway becomes important later in the stress, taken in account that drought stress is not yet present at 24h. Especially considering that although the *scl14* mutant line showed a fitness comparable to WT under drought stress (data not shown), it is part of a multigenetic family: 33 GRAS proteins are encoded by Arabidopsis genome ^40^. Furthermore, at least two other members are present in the more stringent LISCL class of GRAS proteins: AtSCL9 and AtSCL11 ^41^, and AtSCL33 was suggested to be a paralogue of SCL14 ^39^. On the other hand, the amazing fitness of OE:SCL14 plants both under control conditions and when treated with β-CCA highlights the role of SCL14-dependent cellular detoxification in plant protection from drought stress. Interestingly, rice plants overexpressing the OsGRAS23 protein, an homolog of AtSCL14, AtSCL11 and AtSCL9, have already been reported as resistant to drought stress ^41^, supporting our results in Arabidopsis, and further encouraging the use of β-CCA on these genotypes.

**Fig. 6.**
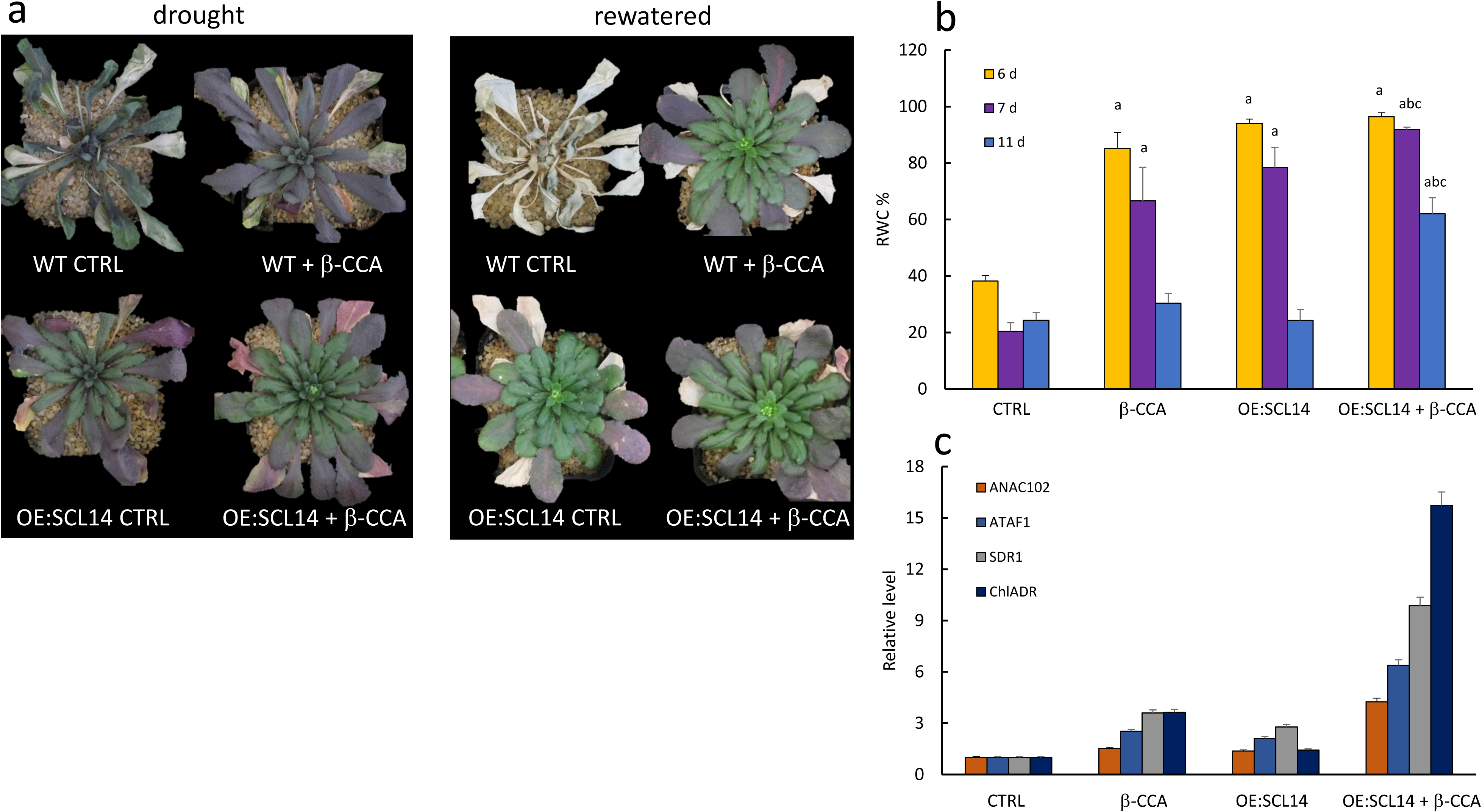
The β-CCA protective effect was enhanced by overexpressing SCL14. a) picture of the plants (WT and OE:SCL14) pre-treated with β-CCA or with water (CTRL) and exposed to water stress for 11 days and after rewatering (2 plants). b) RWC of WT and OE:SCL14 leaves after 6, 8 or 11 days of water stress (2 plants, mean values of 6 leaves per point). c) Expression levels of marker genes for the SCL14 response (3 plants pooled, 4 technical replicates).

Conversely to β-CCA, pre-exposure of plants for 4 h to an atmosphere containing volatile β-CC lead to a strong upregulation of the SCL14-dependent response ^38^. Similar to OE:SCL14 plants, the treatment with β-CC led to an enhancement of drought tolerance (SI Appendix, Fig. S5), as did β-CCA. We can, therefore, hypothesize that the strong induction of the SCL14 pathway by β-CC, together with the generation of β-CCA, could lead to an even stronger response than β-CCA by itself, although the volatile nature of β-CC could compromise its use in the field. These results confirm that the acclimatory response triggered by β-CCA is only part of the response induced by β-CC, while β-CC triggers the β-CCA response indeed.

In summary, we have identified a new signaling molecule in plants, β-CCA, downstream of the apocarotenoid β-CC, which triggers the tolerance of plants towards drought stress by activating a branch of the β-CC-dependent signaling mechanism. This work provides thus a new protective agent that could be exploited to increase plant tolerance to drought using simple application procedures. In addition, the use of β-CCA in the SCL14 or OsGRAS23-overexpressing plants could further boost the protective effect. In any case, additional studies will have to identify key components of the drought defense mechanism proper to β-CCA, and a genetic screen is in progress to isolate β-CCA-insensitive Arabidopsis mutants.

Although the molecular mechanism elicited by β-CCA is still elusive, it is conserved in several plant species such as pepper (*Capsicum*), pansy flower plants (*Viola tricolor*) (SI Appendix, Fig. S3) and tomato (*Solanum lycopersicum*). In fact, we have exposed tomato plants to a rather moderate drought stress in outdoor experiments to illustrate the potential of applications of this compound (Fig. 7). Similar to Arabidopsis, leaves of β-CCA treated tomato plants retained more water (Fig. 7*A*) and showed less symptoms of leaf dehydration during drought stress (Fig. 7*B*) than untreated plants. After recovery from the water stress period, tomato fruit size of β-CCA-treated plants was noticeably enhanced compared to fruits of control plants: the fruit size and weight were around 30% higher (Fig. 7C and D, SI Appendix Fig. S6), while the number of tomatoes was not affected (Fig. 7*E*). The experiment of Fig. 7 shows that the protective effect of β-CCA takes place also under outdoor conditions when drought is combined with temperature and light changes and can have marked beneficial effects on plant productivity. Furthermore, similarly to what we observed with Arabidopsis, leaves of tomato plants irrigated with β-CCA-containing water accumulated high amounts of β-CCA (Fig. 7*F*). Interestingly, this accumulation was not found in the fruit flesh, which contained similar β-CCA levels in treated and untreated plants (Fig. 7*G*).

**Fig. 7.**
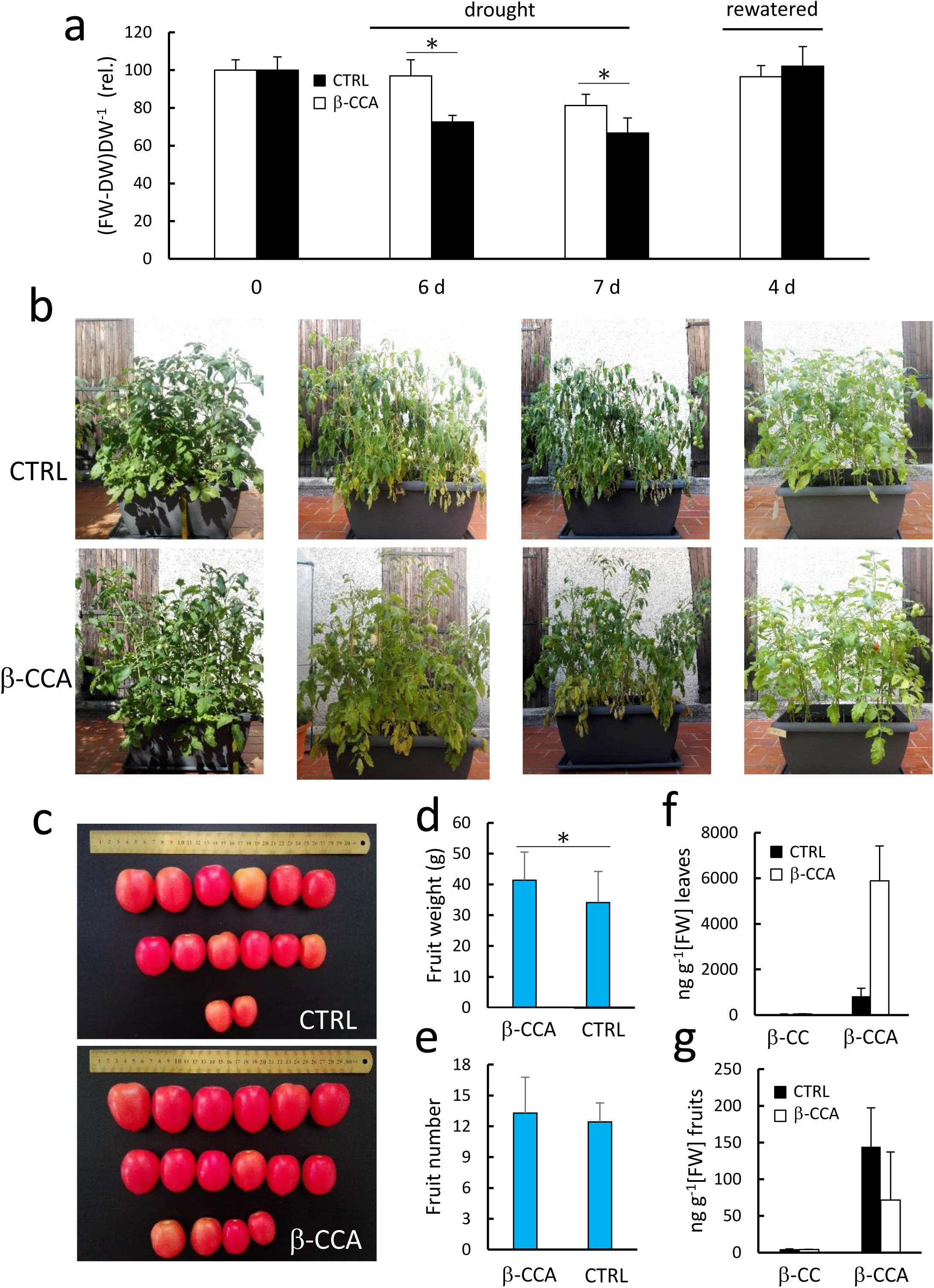
β-CCA protects tomato plants during water stress in outdoor experiments. a) Leaf water content of tomato plants (pre-treated with β-CCA or with water (CTRL)) during water stress (6 d and 7 d after stopping watering) and recovery (4 d after re-watering) (8 plants, mean values of 8 leaves per point). b) Picture of the plants pre-treated with β-CCA or with water (CTRL) and exposed to water stress for 6 d and 7 d (8 plants). c) Tomato fruits from the first and second truss were harvested when ripe, from plants treated or untreated with β-CCA and exposed to water stress for 8 d (30 fruits). d) Average weight of the harvested tomato fruits (30 fruits). e) Average number of fruits per plant (8 plants). f) and g) β-CC and β-CCA levels in tomato leaves (3 leaves, 2 technical replicates) (f) and in tomato fruits (3 fruits, 3 technical replicates) (g). *, different at P <0.01 (Student’s t-test).

Previous attempts to enhance plant tolerance to drought by application of exogenous compounds essentially concern the apocarotenoid phytohormone abscisic acid (ABA) ^42,43^ or its synthetic agonists, quinabactin ^44^, which can reduce transpirational water losses through stomatal closure ^27^. β-CCA has a different mode of action (Fig. 2 and Fig. 3) and possesses a number of advantages over those anti-transpirants, including low cost, stability and protective action independent of stomatal regulation, thus limiting the effects on carbon acquisition. In relation with the latter feature, 3-week application of β-CCA was found to have no inhibitory effect on plant growth (SI Appendix, Fig. S7), indicating that this compound is not harmful to plants in the long term, at the applied concentrations. This is at variance with ABA which was reported to reduce the growth rate of shoots both under osmotic stress and under normal conditions ^45–47^. Acetate is another metabolite that was recently shown to provide some protection against drought stress ^31^. However, its phytotoxicity is well known and is even exploited in the elaboration of herbicides ^48^, and this may limit its applicability as a phytoprotector ^48^. Independently of the molecular mechanisms that remains to be identified, the β-CCA effect on drought tolerance, reported here, show that this carotenoid-derived compound has great potential for agronomical applications in the protection of crops as an anti-drought agent.

## Materials and Methods

### Plant growth and stress treatment

Wild type (WT, ecotype Col 0), *ost2-2, mbs1, scl14* and *abi1* mutant *Arabidopsis thaliana* lines and the *SCL14*-overexpressing line (OE:SCL14) were grown for 5 weeks in short-day conditions (8h/16h, day/night) under a moderate photon flux density (PFD) of ∼150 µmol photons m^-2^ s^-1^, controlled temperature (20 °C/18 °C, day/night) under a relative air humidity of 65 %. For the drought experiments, Arabidopsis plants were watered with 25 ml per pot of water acidified with citric acid (1.5 mM) or of water containing 1.5 mM β-CCA. Water stress was subsequently applied by stopping watering. Pepper and pansy plants were bought on the local market and acclimated in a greenhouse for 1 week. Drought stress was applied by stopping watering after the treatment with 500 ml water or 0.15 mM β-CCA per pot. Tomato plants (*Solanum lycopersicum,* cultivar Rio grande) were grown in 80-L pots (8 plants per pot). When fruits in first truss were mature and started changing their color, drought stress was applied by stopping watering after the treatment with 4 L water or 1.5 mM β-CCA per pot. Treatments of Arabidopsis plants with volatile β-CC in transparent airtight plexiglass box were performed as previously explained (1).

### β-cyclocitral oxidation

1 ml of pure β-cyclocitral (Sigma-Aldrich) was transferred in 1 L of bi-distilled water, in a closed container, and agitated for 1 d. Then the container was opened and agitated for 1 additional day. 50 μl samples of the solution were taken at different times for β-CCA analyses.

### GC/MS measurements

GC/MS analyses were performed as described previously (1). The lipid fraction was extracted from the samples (aqueous solutions or about 500 mg plant tissues) in 4 ml *tert*-Butyl methyl ether plus 1 ml 1mM HCl, containing 4-nonanol as internal reference (10 μg). After centrifugation, the supernatant was collected, evaporated and analyzed by GC-MS. β-CC and β-CCA were quantified on the most probable ion (m/z 137 and 153, respectively) in SIM analyses. The mass spectrum of β-CCA can be found in (13). Pure β-CCA can be produced as reported in (13).

### Stomatal conductance measurements

Stomatal conductance was analyzed on at least two leaves per plants on six plants per condition, using an AP4 diffusive porometer (Delta-T Devices, Cambridge, UK). Two readings were taken per leaves and averaged. Measurements were made on the abaxial leaf surface between 1 h and 2 h after the start of the illumination. Stomatal conductance measurement by IRGA were performed with a LI-COR 6400 (LiCor Inc., Lincoln, NE, USA) equipped with a clamp-on leaf cuvette (6400-40 Leaf Chamber Fluorometer; LiCor Inc.). Leaf temperature was maintained at 22 °C, and leaf-to-air VPD was at 2 kPa. A 10% blue and 90% red light was provided from a LED array on top-side cuvette and set to 500 µmol m^-2^ s^-1^. Stomatal conductance was analyzed on one leaf per plant on six plants per condition, after 30 minutes of equilibration.

### Relative water content (RWC)

Six leaf disks per condition were cut from leaves of β-CCA-treated or -untreated plants and weighted (FW). For measuring the turgid weight (TW), samples were submerged with bi-distilled water and left at 4°C for 16 hours. Dry weight (DW) was measured after 16h at 70°C in a ventilated oven. RWC was measured following the formula (FW-DW)/(TW-DW) x 100.

### Ion Leakage

Cell membrane stability can be estimated by the use of electrolyte leakage method (43). Six leaves per condition were cut from β-CCA-treated or -untreated plants and weighted (FW), and placed in 25 ml of bi-distilled water. Conductivity of the solution was measured with a DIST-5 Hanna conductometer after 2 h of mild agitation and after boiling the samples. The values were normalized on the FW and the values after boiling are reported as the maximal value.

### Leaf osmotic potential

Leaf osmotic potential was measured on freeze-thawed leaf discs using C-52 psychrometer chambers and Psypro control unit (Wescor, Logan, UT, USA). Leaf discs of 7 mm diameter were placed in sealed plastic tubes, frozen in liquid nitrogen and kept at −80 °C. For measurements, the discs were defrosted for 2 min and transferred onto sample holders of psychrometer chambers. To speed up the equilibration time of water potential in the chambers, leaf discs were pierced several times before enclosure (44).

### RNA isolation and qRT-PCR

Total RNA was isolated from 100 mg leaves using the Nucleospin^®^ RNA Plant kit (Macherey-Nagel). The concentration was measured on a NanoDrop2000 (Thermo Scientific, USA). First strand cDNA was synthesized from 1 µg total RNA using the PrimeScript™ Reverse Transcriptase kit (Takara, Japan). qRT-PCR was performed on a Lightcycler 480 Real-Time PCR system (Roche, Switzerland). 3 µl of a reaction mixture comprising SYBR Green I Master (Roche, Switzerland), 10 µM each of forward and reverse primers and water, was added to 2 µL of a 10-fold diluted cDNA sample in a 384 well plate. The PCR program used was: 95 °C for 10 min, then 45 cycles of 95 °C for 15 s, 58 °C for 15 s and 72 °C for 15 s. At least three biological replicates were performed for each gene tested. Primers for all genes examined (SI Appendix, Table S1) were designed using the Primer3Plus software. Profilin-1 (PRF1, AT2G19760) and Cyclophilin 5 (CYP5, AT2G29960) were used as reference genes for the normalization of gene expression levels.

### Statistical analysis

All experiments were repeated at least two times, and the images represent typical examples. The values are represented as the means + standard deviation. The statistical significance was tested using Student’s t-test (two-tailed, unequal variances). Sample size is reported in figure legends as number of plants per experiment.

## Supporting information

## ACKNOWLEDGMENTS

We would like to thank the GRAP platform (CEA/Cadarache) for growing plants under normal and stress conditions. We also thank Pierre Chagvardieff (CEA/Cadarache) for support and Giovanna Loro for her assistance. The *mbs1* mutant was received from Ralf Bock (Max Planck Institute, Golm, Germany). The *scl14* and OE:SCL14 were a kind gift of Christiane Gatz (Gottingen, Germany). This work was funded by the French National Research Agency (ANR project SLOSAM, 14-CE02-0010-02) (M.H.). Y.M. was supported by the European Union (project WATBIO, grant agreement no. 311929) and the French National Research Agency (ANR, project ORCA, grant agreement no. ANR-13-BS06-0005-01). CEA has submitted a provisional patent on behalf of S.D.A. and M.H. on aspects of the findings.

