## Supplementary material for "β-cyclocitric acid: a new apocarotenoid eliciting drought tolerance in plants"

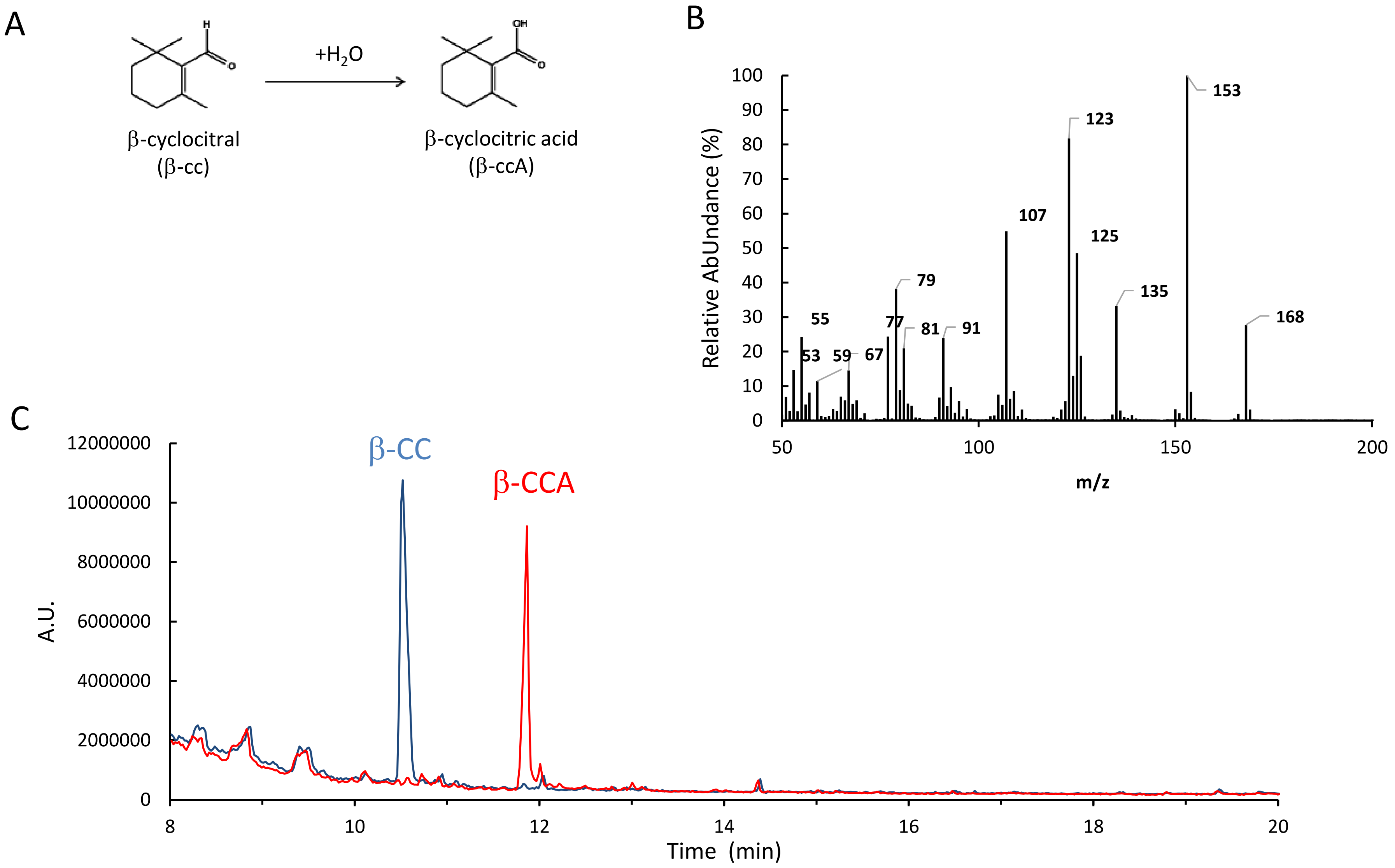


**Supplementary Fig. S1**. (A) β-cyclocitral (β-CC) oxidizes to β-cyclocitric acid (β-CCA) in water. (B) Fragmentation plot of the peak corresponding to β-CCA. (C) GC-MS analysis of the oxidation of β-CC (m/z 137) into β-CCA (m/z 153). β-CC was injected in water and mixed in a closed bottle for 24 h and in an open bottle for 3 d. The graph shows GC-MS scan (TIC) at time 0 (in blue) and at time 4 d (in red). After 4 d, β-CCA is the only remarkable species in the solution with a concentration of about 1.5 mM.

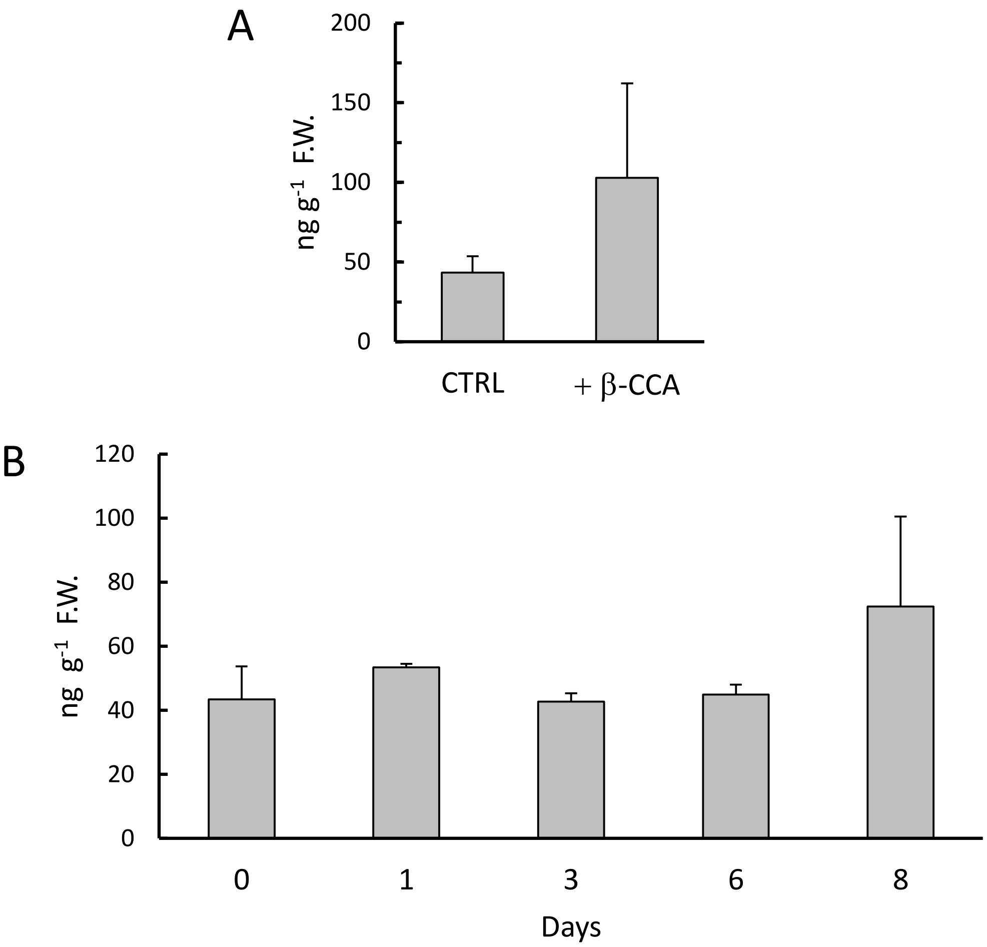


**Supplementary Fig. S2**. β-CC levels in leaves treated with β-CCA. A) Leaves directly treated with β-CCA (spray) (experiment of Fig. 1d). B) Leaves from plants watered with β-CCA (experiment of Fig. 1e).

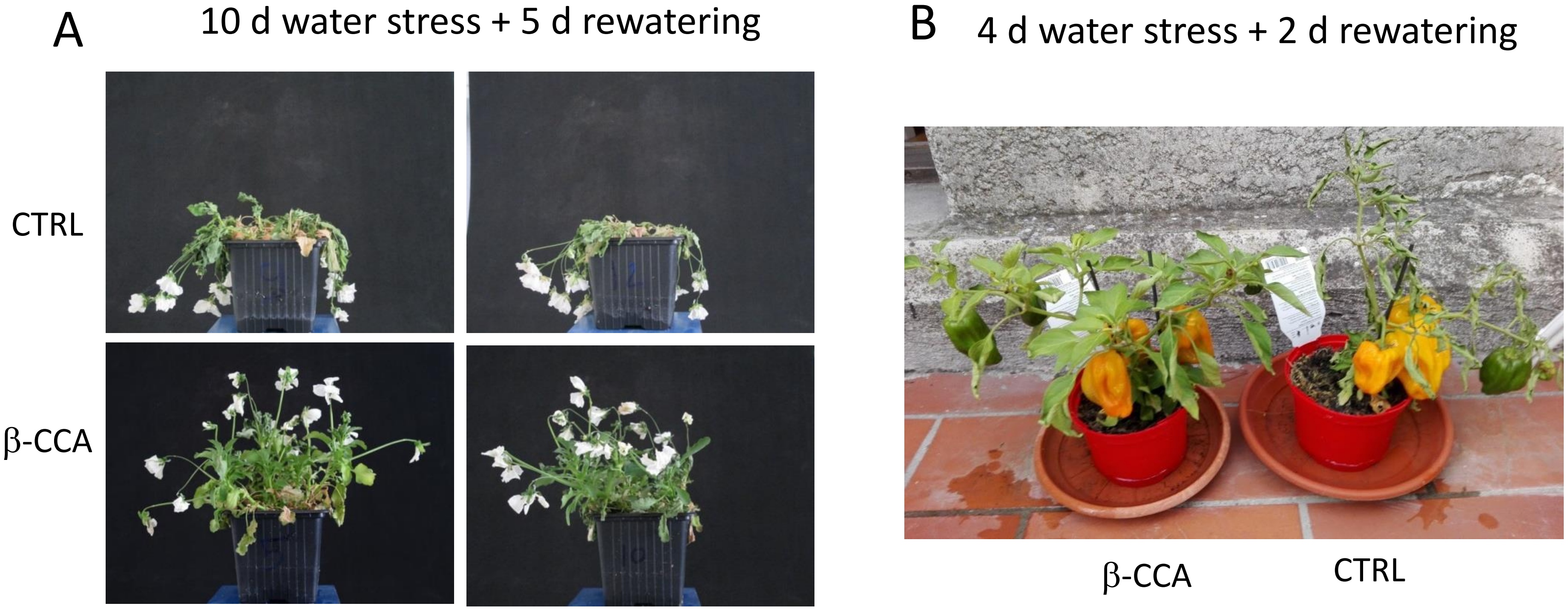


**Supplementary Fig. S3**. Protective effect of β-CCA against drought stress in different plant species. A) Pansy flower plants (*Viola bicolor*) exposed for 10 d to water stress by withdrawing watering followed by rewatering for 5 d. b) Pepper plants (*Capsicum annuum*) exposed for 4 d to water stress by withdrawing watering followed by 2 d of rewatering.

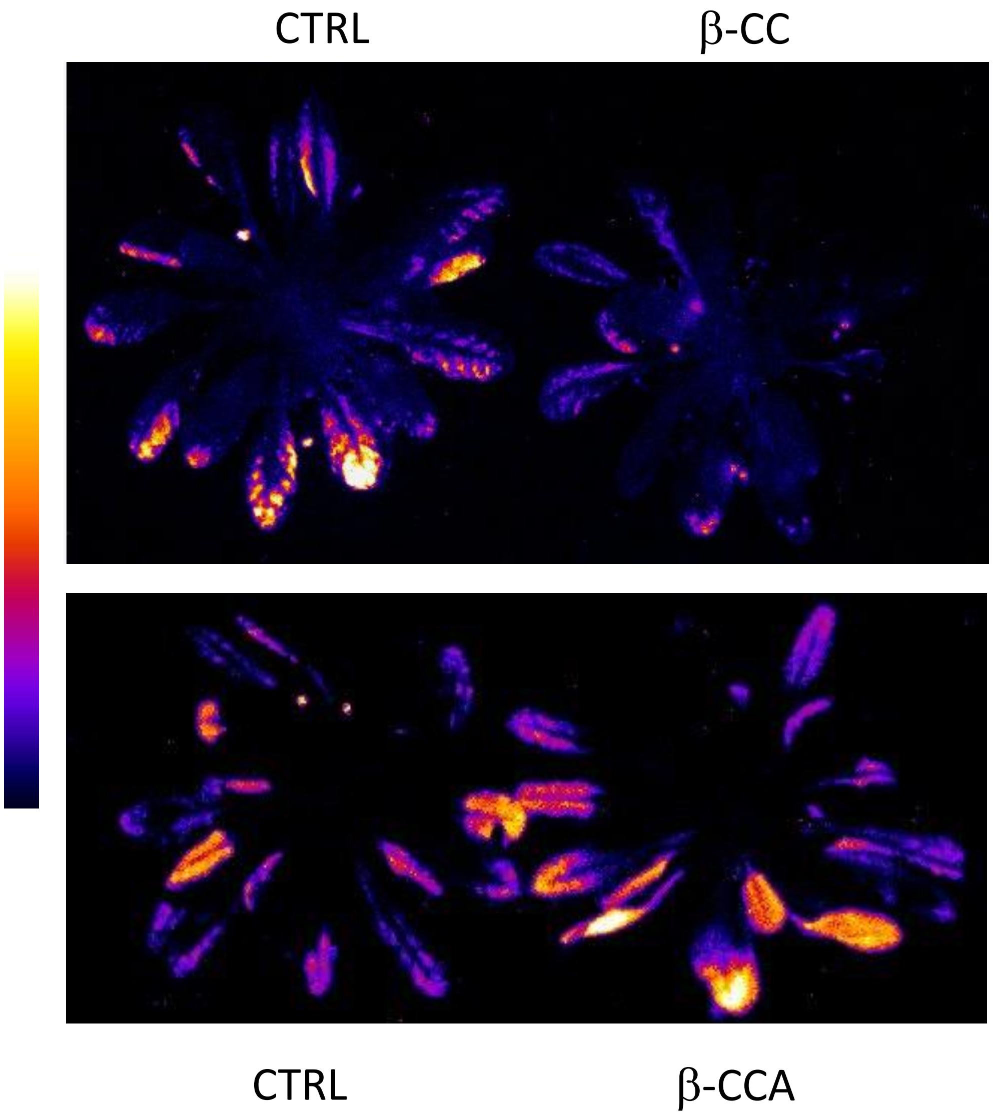


**Supplementary Fig. S4**. Effect of high light (1500 μmol photons $m^{-2}$ $s^{-1}$) at low temperature (7°C) for 24 h on control (CTRL), β-CC-pretreated and β-CCA-pretreated Arabidopsis plants. Plants were exposed to volatile β-CC for 4 h. For the β-CCA treatment, plants were exposed high light and cold 48 h after watering the plants with 2.5 mM β-CCA-containing water. Lipid peroxidation was measured by autoluminescence imaging.

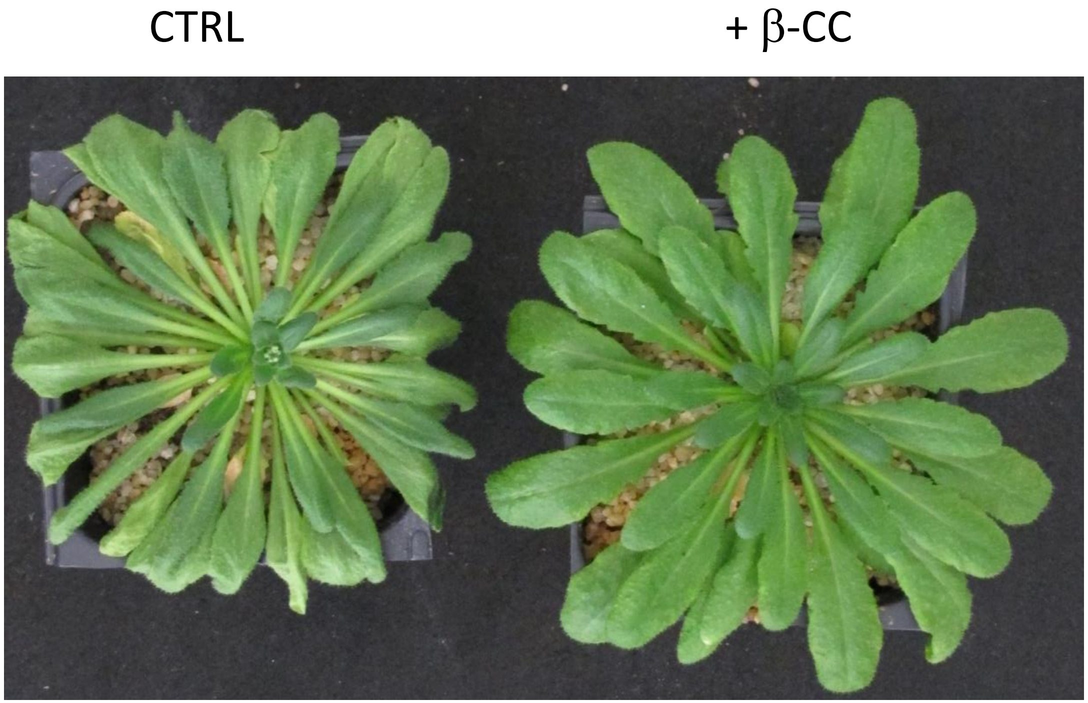


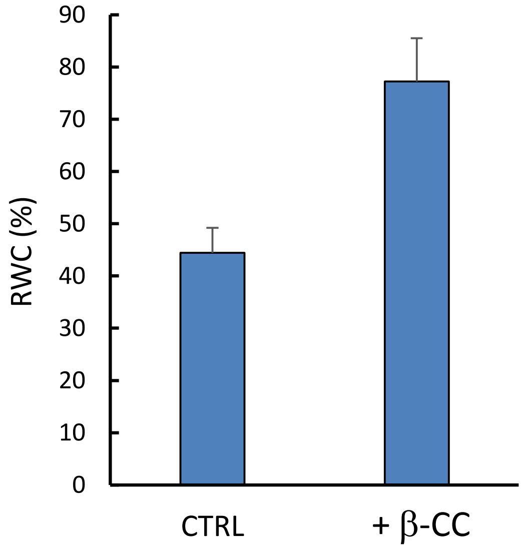


**Supplementary Fig. S5**. Effect of β-CC on the tolerance of Arabidopsis plants to drought stress. Plants were pre-exposed for 4 h to volatile β-CC (100 µl) in closed chambers, as described in Ramel et al. (2012). Control plants were exposed to the same conditions with β-CC being replaced by water. Drought stress was subsequently induced by stopping watering for 6 d. A) Effect of water deprivation on control (crtl) and β-CC-pretreated plants. B) RWC of the water-stressed plants.

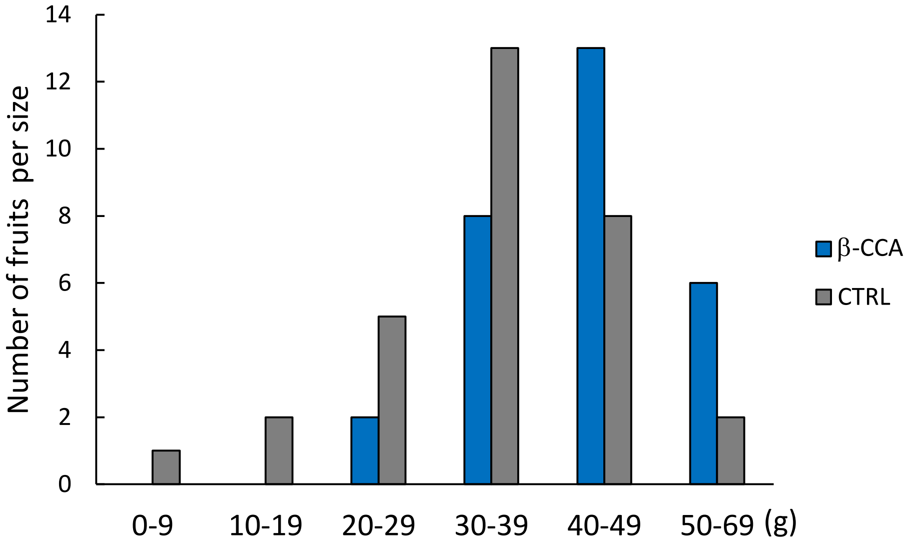


**Supplementary Fig. S6**. Number of fruit per size class (g) of the experiment used for calculating the average weight of fruits (Figure 7 D)

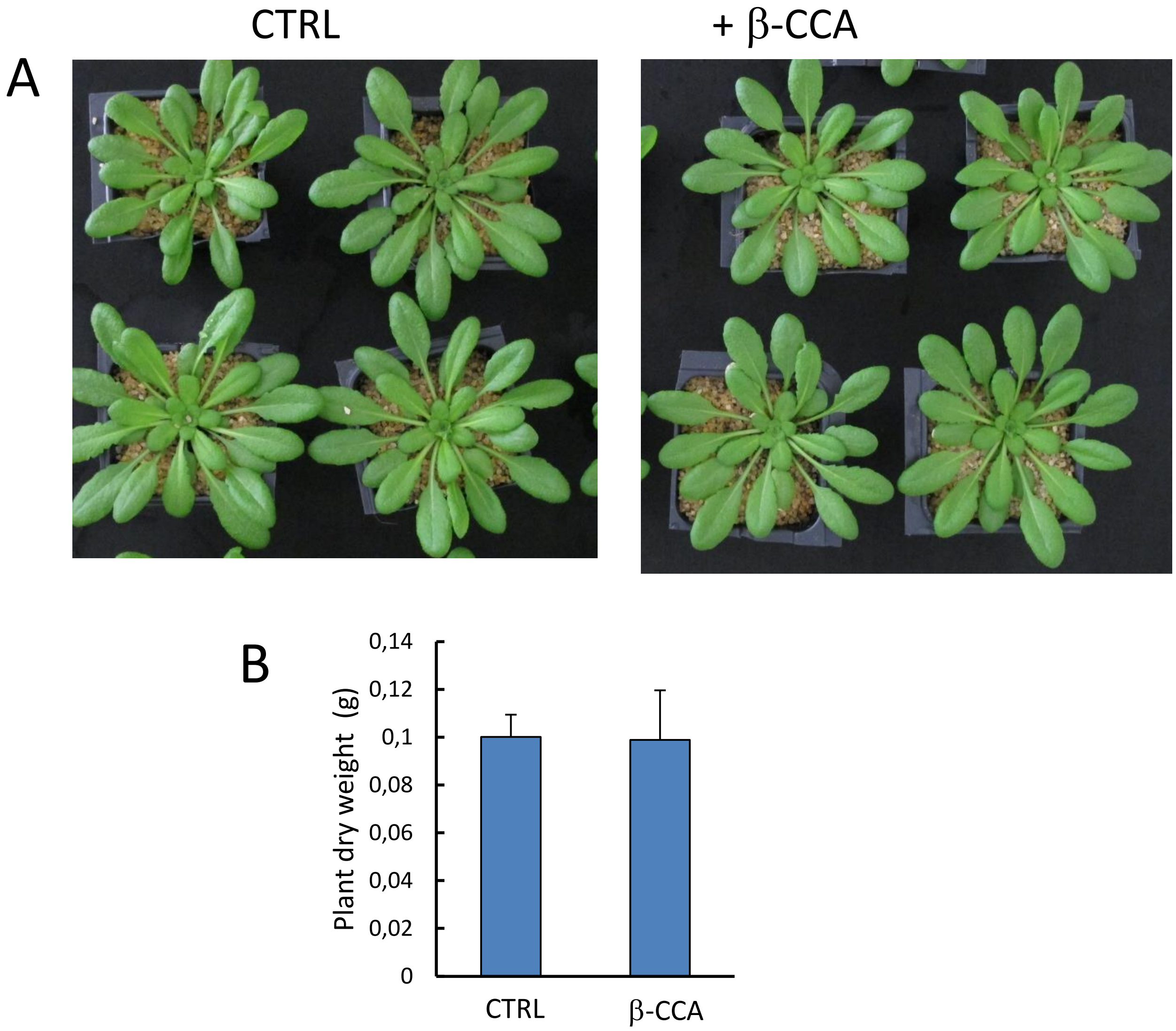


**Supplementary Fig. S7**. Long-term exposure of Arabidopsis plants to β-CCA had no effect on the growth rate. Plants aged 10 d were watered for 30 d with water containing β-CCA in order to provide plants with 6 mg β-CCA per plant and per week. A) picture of the plants and B) shoot dry weight at the end of the experiment.
